# Constructing and analysing dynamic models with modelbase v1.0 - a software update

**DOI:** 10.1101/2020.09.30.321380

**Authors:** Marvin van Aalst, Oliver Ebenhöh, Anna Matuszyńska

## Abstract

**Background:** Computational mathematical models of biological and biomedical systems have been successfully applied to advance our understanding of various regulatory processes, metabolic fluxes, effects of drug therapies and disease evolution or transmission. Unfortunately, despite community efforts leading to the development of SBML or the BioModels database, many published models have not been fully exploited, largely due to lack of proper documentation or the dependence on proprietary software. To facilitate synergies within the emerging research fields of systems biology and medicine by reusing and further developing existing models, an open-source toolbox that makes the overall process of model construction more consistent, understandable, transparent and reproducible is desired.

**Results and Discussion:** We provide here the update on the development of `modelbase`, a free expandable Python package for constructing and analysing ordinary differential equation-based mathematical models of dynamic systems. It provides intuitive and unified methods to construct and solve these systems. Significantly expanded visualisation methods allow convenient analyses of structural and dynamic properties of the models. Specifying reaction stoichiometries and rate equations, the system of differential equations is assembled automatically. A newly provided library of common kinetic rate laws highly reduces the repetitiveness of the computer programming code, and provides full SBML compatibility. Previous versions provided functions for automatic construction of networks for isotope labelling studies. Using user-provided label maps, `modelbase` v1.0 streamlines the expansion of classic models to their isotope-specific versions. Finally, the library of previously published models implemented in `modelbase` is continuously growing. Ranging from photosynthesis over tumour cell growth to viral infection evolution, all models are available now in a transparent, reusable and unified format using `modelbase`.

**Conclusion:** With the small price of learning a new software package, which is written in Python, currently one of the most popular programming languages, the user can develop new models and actively profit from the work of others, repeating and reproducing models in a consistent, tractable and expandable manner. Moreover, the expansion of models to their label specific versions enables simulating label propagation, thus providing quantitative information regarding network topology and metabolic fluxes.

## Background

Mathematical models are accepted as valuable tools in advancing biological and medical research [1,2]. In particular, models based on ordinary differential equations (ODE) found their application in a variety of fields. Most recently, deterministic models simulating the dynamics of infectious diseases gained the interest of the general public during our combat of the Covid-19 pandemic, when a large number of ODE based mathematical models has been developed and discussed even in nonscientific journals (see for example [3–5]). Such focus on mathematical modelling is not surprising, because computational models allow for methodical investigations of complex systems under fixed, controlled and reproducible conditions. Hence, the effect of various perturbations of the systems *in silico* can be inspected systematically. Importantly, long before exploring their predictive power, the model building process itself plays an important role in integrating and systematising vast amounts of available information [6]. Properly designed and verified computational models can be used to develop hypotheses to guide the design of new research experiments (e.g., in immunology to study lymphoid tissue formation [7]), support metabolic engineering efforts (e.g., identification of enzymes to enhance essential oil production in peppermint [8]), contribute to tailoring medical treatment to the individual patient in the spirit of precision medicine (e.g., in oncology [2]), or guide political decision making and governmental strategies (see the review on the impact of modelling for European Union Policy [9]). Considering their potential impact, it is crucial that models are openly accessible so that they can be verified and corrected, if necessary.

In many publications, modelling efforts are justified by the emergence of extraordinary amounts of data provided by new experimental techniques. However, arguing for the necessity of model construction only because a certain type or amount of data exists, ignores several important aspects. Firstly, computational models are generally a result of months, if not years, of intense research, which involves gathering and sorting information, simplifying numerous details and distilling out the essentials, implementing the mathematical description in computer code, carrying out performance tests and, finally, validation of the simulation results. Our understanding of many phenomena could become deeper if instead of constructing yet another first-generation model, we could efficiently build on the knowledge that was systematically collected in previously developed models. Secondly, the invaluable knowledge generated during the model construction process is often lost, mainly because of the main developer leaves the research team, but also due to unfavourable funding strategies. It is easier to obtain research funds for the construction of novel, even if perfunctory models, than to support a long-term maintenance of existing ones. Preservation of the information collected in the form of a computational model has became an important quest in systems biology, and has been to some extend addressed by the community. Development of the Systems Biology Markup Language (SBML) [10] for unified communication and storage of biomedical computational models and the existence of the BioModels repository [11] already ensured the survival of constructed models beyond the academic lifetime of their developers or the lifetime of the software used to create them. But a completed model in the SBML format does not allow to follow the logic of model construction and the knowledge generated by the building process is not preserved. Such knowledge loss can be prevented by providing simple-to-use toolboxes enforcing a universal and readable form of constructing models.

We have therefore decided to develop `modelbase` [12], a Python package that encourages the user to actively engage in the model building process. On the one hand we fix the core of the model construction process, while on the other hand the software does not make the definitions too strict, and fully integrates the model construction process into the Python programming language. This differentiates our software from many available Python-based modelling tools (such as ScrumPy [13] or PySCeS [14]) and other mathematical modelling languages (recently reviewed from a software engineering perspective by Schölzel and colleagues [15]). We report here new features in `modelbase` v1.0, developed over the last two years. We have significantly improved the interface to make model construction easier and more intuitive. The accompanying repository of re-implemented, published models has been considerably expanded, and now includes a diverse selection of biomedical models. This diversity highlights the general applicability of our software. Essentially, every dynamic process that can be described by a system of ODEs can be implemented with `modelbase`.

## Implementation

`modelbase` is a Python package to facilitate construction and analysis of ODE based mathematical models of biological systems. Version 1.0 introduces changes not compatible with the previous official release 0.2.5 published in [12]. All API changes are summarised in the official documentation hosted by ReadTheDocs.

The model building process starts by creating a modelling object in the dedicated Python class `Model` and adding to it the chemical compounds of the system. Then, following the intuition of connecting the compounds, you construct the network by adding the reactions one by one. Each reaction requires stoichiometric coefficients and a kinetic rate law. The latter can be provided either as a custom function or selecting from the newly provided library of rate laws. The usage of this library (`ratelaws`) reduces the repetitiveness by avoiding boilerplate code. It requires the user to explicitly define reaction properties, such as directionality. This contributes to a systematic and understandable construction process, following the second guideline from the Zen of Python: “Explicit is better than implicit”.

From this, `modelbase` automatically assembles the system of ODEs. It also provides numerous methods to conveniently retrieve information about the constructed model. In particular, the `get_*` methods allow inspecting all components of the model, and calculate reaction rates for given concentration values. These functions have multiple variants to return all common data structures (array, dictionary, data frames).

After the model building process is completed, simulation and analyses of the model are performed with the `Simulator` class. Currently, we offer interfaces to two integrators to solve stiff and non-stiff ODE systems. Provided you have installed the Assimulo package [16], as recommended in our installation guide, `modelbase` will be using CVode, a variable-order, variable-step multi-step algorithm. The CVode class provides a direct connection to Sundials, the SUite of Nonlinear and DIfferential/ALgebraic equation Solvers [17] which is a powerful industrial solver and robust time integrator, with a high computing performance. In case when Assimulo is not available, the software will automatically switch to the SciPy library [18] using lsoda as an integrator, which in our experience showed a lower computing performance.

### Metabolic Control Analysis

Sensitivity analysis provides a theoretical foundation to systematically quantify effects of small parameter perturbations on the global system behaviour. In particular, Metabolic Control Analysis (MCA), initially developed to study metabolic systems, is an important and widely used framework providing quantitative information about the response of the system to perturbations [19, 20]. This new version of `modelbase` now has a full suite of methods to calculate response coefficients and elasticises, and plotting them as a heat-map, giving a clear and intuitive colour-coded visualisation of the results.

An example of such visualisation, for a re-implemented toy model of the upper part of glycolysis (Section 3.1.2 [21]), can be found in Figure 1.

**Figure 1.**
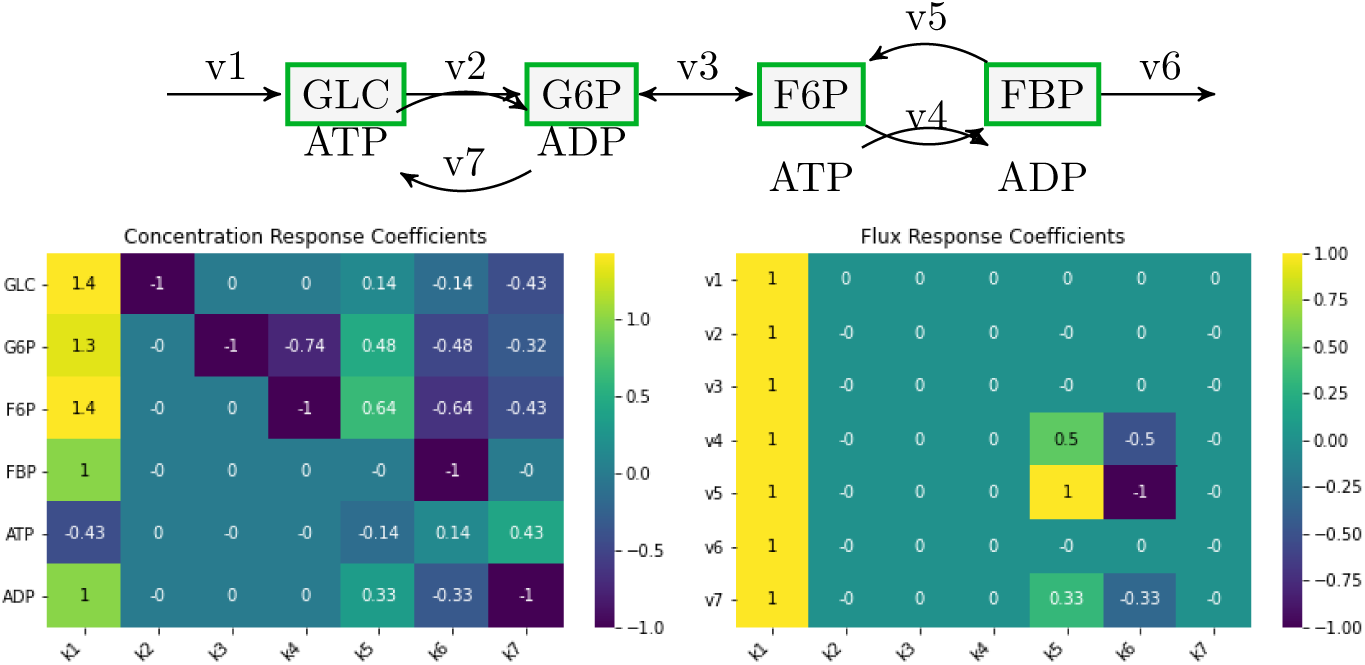
Metabolic Control Analysis of the toy model of the upper glycolysis. Using the mca class we have calculated the concentration (left) and flux (right) control coefficients for a toy model of upper glycolysis introduced for educational purpose in [21], and visualised them in a form of a heat map. To reproduce these results run the Jupyter Notebook from the Additional file 1.

### Visualisation support

Many of the available relevant software packages for building computational models restrict the users by providing unmodifiable plotting routines with predefined settings that may not suit your personal preferences. With `modelbase` v1.0 we constructed our plotting functions allowing the user to pass optional keyword-arguments (often abbreviated as **kwargs), so you still access and change all plot elements, providing a transparent and flexible interface to the commonly used matplotlib library [22]. The easy access functions to visualise the results of simulations were expanded from the previous version. They now include plotting selections of compounds or fluxes, phase-plane analysis and the results of MCA.

### Models for isotope tracing

`modelbase` has additionally been developed to aid the *in silico* analyses of label propagation during isotopic studies. In order to simulate the dynamic distribution of isotopes all possible labelling patterns for all intermediates need to be created. By providing an atom transition map in the form of a list or a tuple, all 2^*N*^ isotope-specific versions of a chemical compound are created automatically, where *N* denotes the number of possibly labelled atoms. Changing the name of previous function carbonmap to labelmap in v1.0 acknowledges the diversity of possible labelling experiments that can be reproduced with models built using our software.

#### Isotope tracing under stationary conditions

Sokol and Portais derived the theory of dynamic label propagation under stationary assumption [23]. In steady state the space of possible solutions is reduced and the labelling dynamics can be represented by a set of linear differential equations. We have used this theory and implemented an additional class `LinearLabelModel` which allows rapid calculation of the label propagation given the steady state concentrations and fluxes of the metabolites [23]. `modelbase` will automatically build the linear label model after providing the label maps. Such a model is provided in Figure 2, where we simulate label propagation in a linear non-reversible pathway, as in Figure 1 in [23]. The linear label models are constructed using `modelbase` rate laws and hence can be fully exported as SBML file.

**Figure 2.**
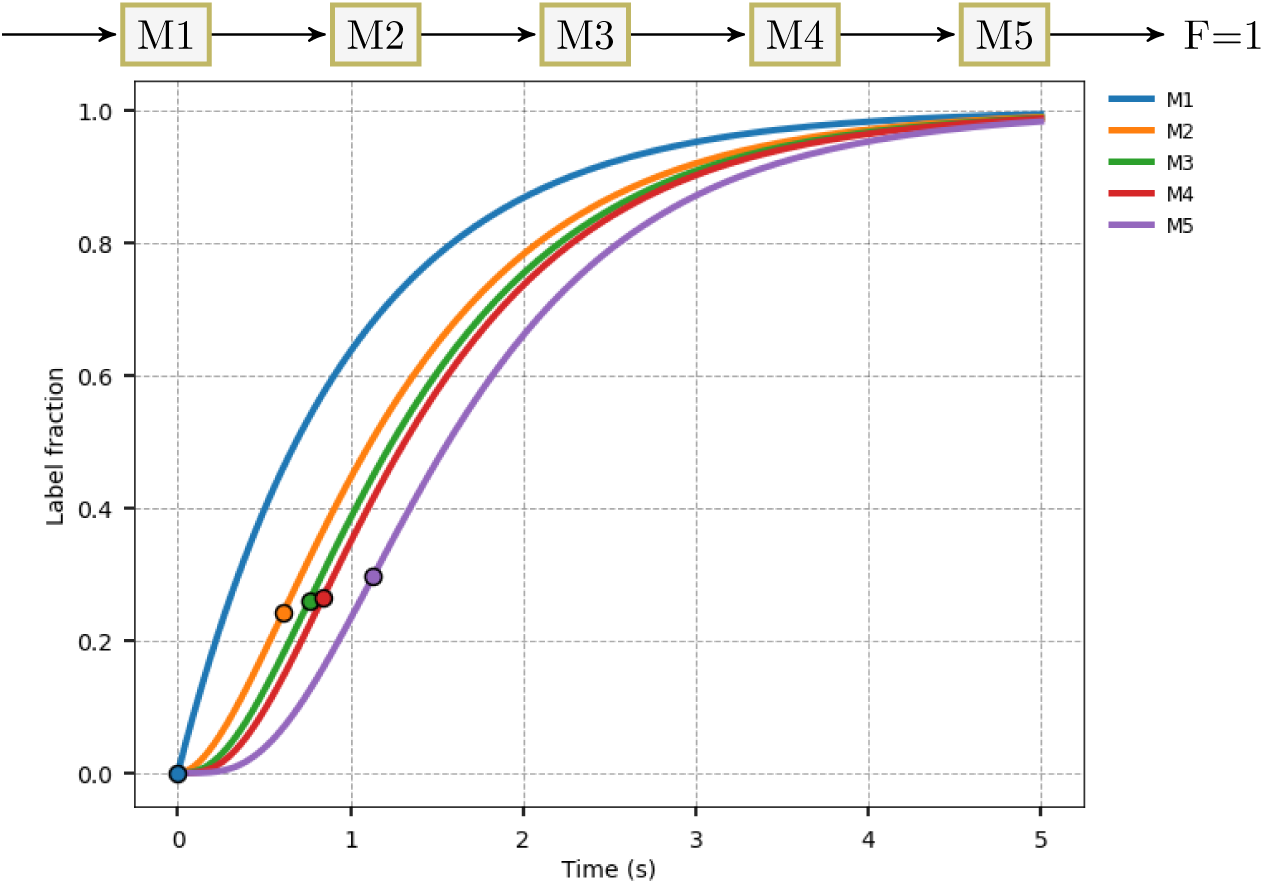
Labelling curves in a linear non-reversible pathway. Example of label propagation curves for a linear non-reversible pathway of 5 randomly sized metabolite pools, as proposed in the paper by Sokol and Portais [23]. Circles mark the position at which the first derivative of each labelling curve reaches maximum. In the original paper this information has been used to analyse the label shock wave (LSW) propagation. To reproduce these results run the Jupyter Notebook from the Additional file 2.

### Model meta data

Many models loose their readability due to the inconsistent, intractable or misguided naming of their components (an example is a model with reactions named as v1 - v10, without referencing them properly). By providing meta data for any `modelbase` object, you can abbreviate component names in a personally meaningful manner and then supply additional annotation information in accordance with standards such as MIRIAM [24] via the newly developed meta data interface. This interface can also be used to supply additional important information compulsorily shared in a publication but not necessarily inside the code, such like the unit of a parameter.

## Results and Discussion

With the newly implemented changes our package becomes more versatile and user friendly. As argued before, its strength lies in its flexibility and applicability to virtually any biological system, with dynamics that can be described using an ODE system. There exist countless mathematical models of biological and biomedical systems derived using ODEs. Many of them are rarely re-used, at least not to the extent that could be reached, if models were shared in a readable, understandable and reusable way [15]. As our package can be efficiently used both for the development of new models, as well as the reconstruction of existing ones, as long as they are published with all kinetic information required for the reconstruction, we hope that `modelbase` will in particular support users with limited modelling experience in re-constructing already existing work, serving as a starting point for their further exploration and development. We have previously demonstrated the versatility of `modelbase` by re-implementing mathematical models previously published without the source code: two models of biochemical processes in plants [25, 26], and a model of the non-oxidative pentose phosphate pathway of human erythrocytes [27, 28]. To present how the software can be applied to study medical systems, we used `modelbase` to re-implement various models, not published by our group, and reproduced key results of the original manuscripts. It was beyond our focus to verify the scientific accuracy of the corresponding model assumptions. We have selected them to show that despite describing different processes, they all share a unified construct. This highlights that by learning how to build a dynamic model with `modelbase`, you in fact do not learn how to build a one-purpose model, but expand your toolbox to be capable of reproducing any given ODE based model. All examples are available as jupyter notebooks and listed in the Additional Files.

### Compartment model for disease evolution

For the purpose of this paper, we surveyed available computational models and subjectively selected a relatively old publication of significant impact, based on the number of citations, published without providing the computational source code, nor details regarding the numerical integration. We have chosen a four compartment model of HIV immunology that investigates the interaction of single virus population with the immune system described only by the CD4^+^ T cells, commonly known as T helper cells [29]. We have implemented the four ODEs describing the dynamics of uninfected (T), latently infected (L) and actively infected CD4^+^ T cells (A) and infectious HIV population (V).

In the Figure 3, we reproduce the results from Fig. 3 from the original paper, where by changing the number of infectious particles produced per actively infected cell (N) we follow the dynamics of the overall T cell population (T+L+A) over a period of 10 years. The model has been further used to explore the effect of azidothymidine, an antiretroviral medication, by decreasing the value of N after 3 years by 25% or 75%, mimicking the blocking of the viral replication of HIV. A more detailed description of the time-dependent drug concentration in the body is often achieved with pharmacokinetic models. Mathematical models based on a system of differential equations that link the dosing regimen with the dynamics of a disease are called pharmacokinetic-pharmacodynamic (PK-PD) models [30] and with the next example we explore how `modelbase` can be used to develop such models.

**Figure 3.**
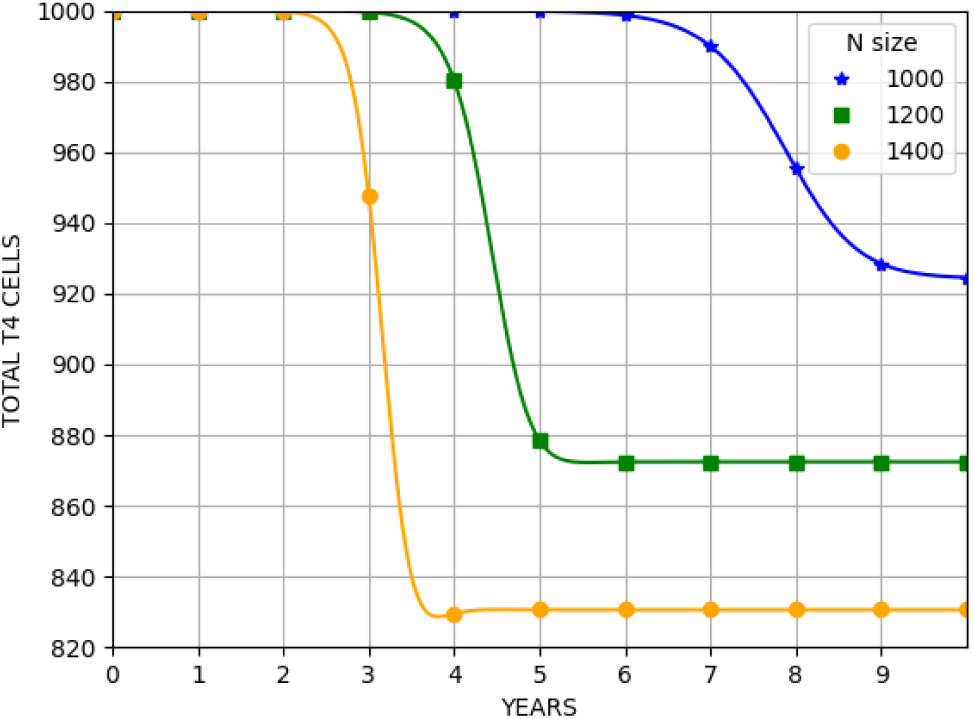
The total CD4+ T-cell population versus time after the infection. We have reproduced the results from Figure 3 from the original paper [29] showing the decrease in the overall population of CD4+ T-cell (uninfected + latently infected + actively infected CD4+) over time, depending on the number of infectious particles produced per actively infected cell (N). To reproduce these results run the Jupyter Notebook from the Additional file 3.

### PK-PD models and precision medicine

The technological advances forced a paradigm shift in many fields of our life, including medicine, making more personalised healthcare not only a possibility but a necessity. A pivotal role in the success of precision medicine will be to correctly determine dosing regimes for drugs [31] and PK-PD models provide a quantitative tool to support this [32]. PK-PD models have proved quite successful in many fields, including oncology and here we used the classical tumour growth model by Simeoni and colleagues, originally implemented using the industry standard software WinNonlin [34]. As the pharmacokinetic model has not been fully described we reproduced only the highly simplified case, where we assume a single drug administration and investigate the effect of drug potency (*k*_2_) on simulated tumour growth curves. In Figure 4 we plot the simulation results of the `modelbase` implementation of the system of four ODEs over the period of 18 days where we systematically changed the value of *k*_2_, assuming a single drug administration on Day 9. With the MCA suite available with our software we can calculate the change of the system in response to perturbation of all other system parameters. Such quantitative description of the systems response to local parameter perturbation provides support in further studies of the rational design of combined drug therapy or discovery of new drug targets, as described in the review by Cascante and colleagues [35].

**Figure 4.**
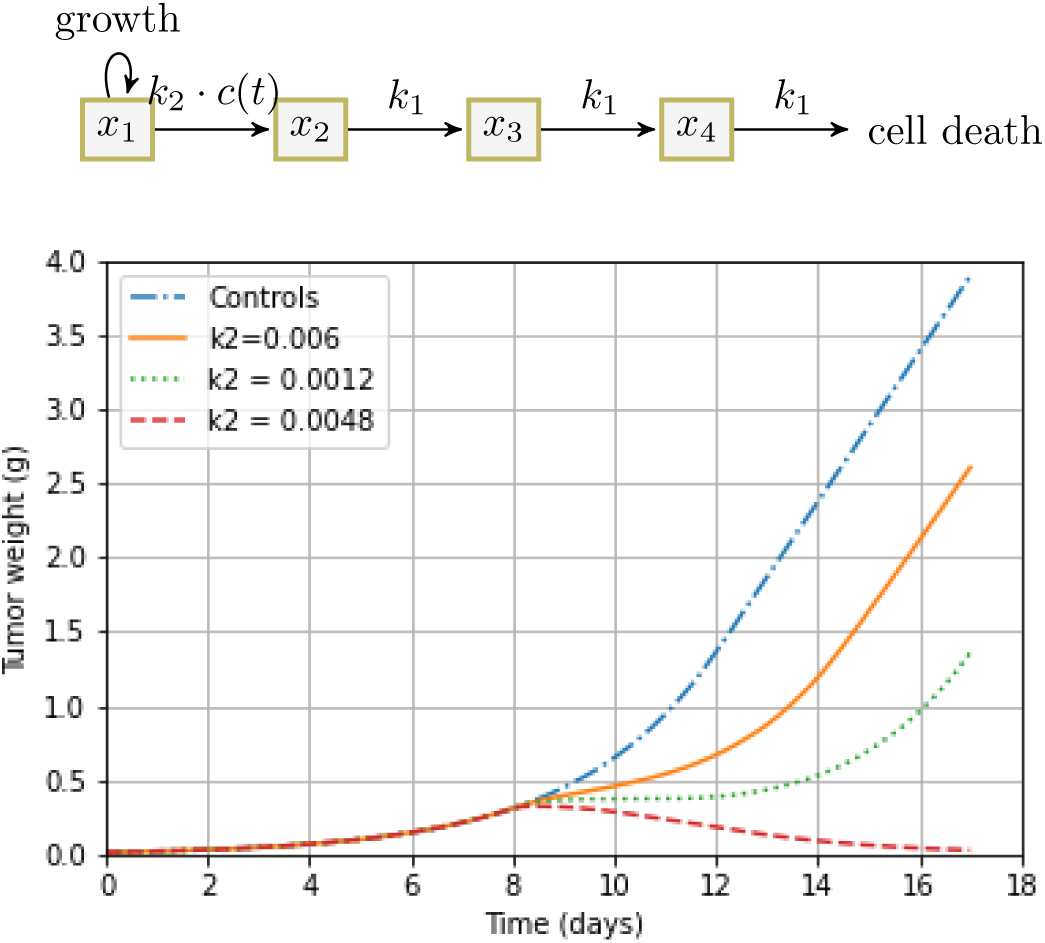
Compartmental pharmacokinetic-pharmacodynamic model of tumour growth after anticancer therapy. We have reproduced the simplified version of the PK-PD model of tumour growth, where PK part is reduced to a single input and simulated the effect of drug potency (*k*_2_) on tumour growth curves. The system of four ODEs describing the dynamics of the system visualised on a scheme above is integrated over the period of 18 days. We systematically changed the value of *k*_2_, assuming a single drug administration on Day 9. We have obtained the same results as in the Figure 3 in the original paper [34]. To reproduce these results run the Jupyter Notebook from the Additional file 4.

### Modelling of infectious diseases with SIR models

Finally, compartmental models based on ODE systems have a long history of application in mathematical epidemiology [36]. Many of them, including numerous recent publications studying the spread of coronavirus, are based on the classic epidemic Susceptible-Infected-Recovered (SIR) model, originating from the theories developed by Kermack and McKendrick at the beginning of last century [37]. One of the most critical information searched for while simulating the dynamics of infectious disease is the existence of disease free or endemic equilibrium and assessment of its stability [38]. Indeed periodic oscillations have been observed for several infectious diseases, including measles, influenza and smallpox [36]. To provide an overview of more `modelbase` functionalities we have implemented a relatively simple SIR model based on the recently published autonomous model for smallpox [39]. We have generated damped oscillations and visualised them using the built-in function `plot_phase_plane` (see Figure 5). In the attached Jupyter notebook we present how quickly and efficiently in terms of lines of code, the SIR model is built and how to add and remove new reactions and/or compounds to construct further variants of this model, such as a SEIR (E-exposed) or SIRD (D-deceased) model.

**Figure 5.**
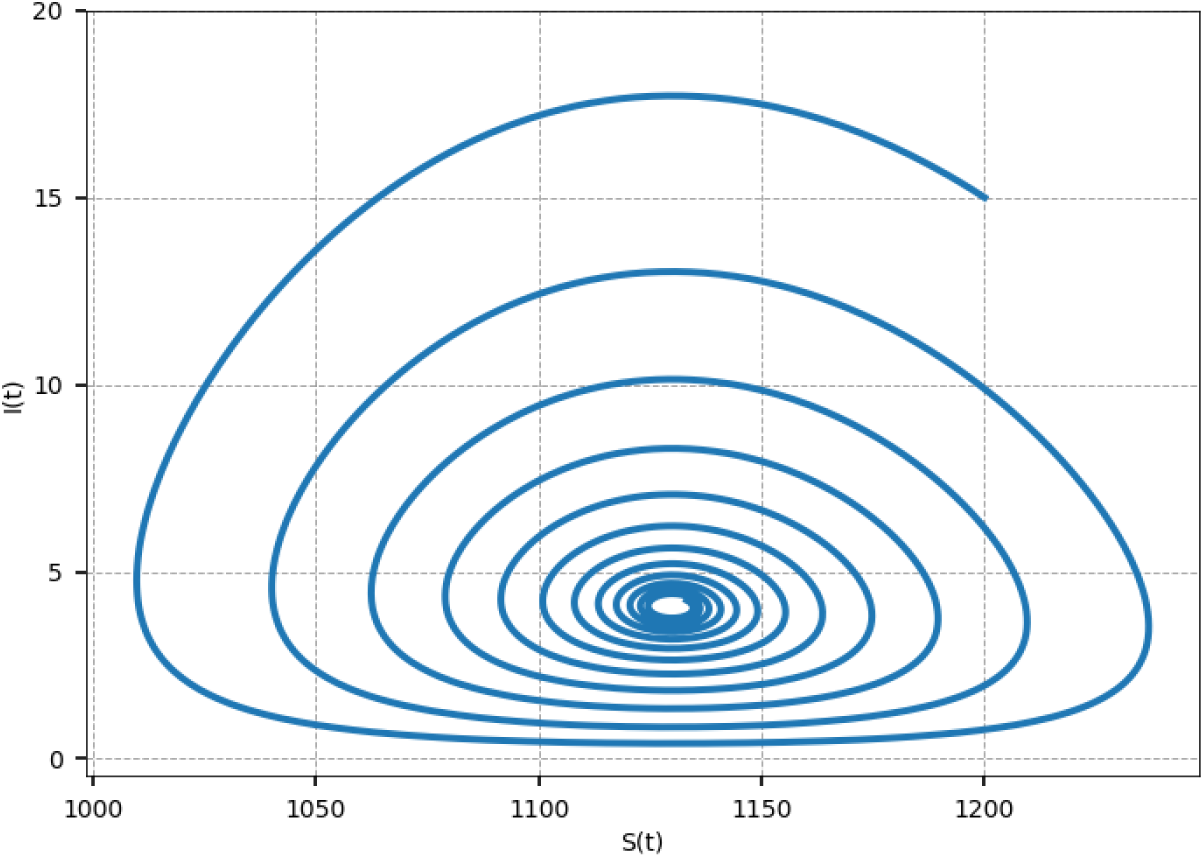
Sample phase portrait obtained with SIR model with oscillations. SIR model with vital dynamics including birth rate has been adapted based on the autonomous model to simulate periodicity of chicken pox outbreak in Hida, Japan [39]. To reproduce these results run the Jupyter Notebook from the Additional file 5.

## Conclusions

We are presenting here updates of our modelling software that has been developed to simplify the building process of mathematical models based on ODEs. `modelbase` is fully embedded in the Python programming language. It facilities a systematic construction of new models, and reproducing models in a consistent, tractable and expandable manner. As ODEs provide a core method to describe the dynamical systems, we hope that our software will serve as the base for deterministic modelling, hence it’s name. With the smoothed interface and clearer description of how the software can be used for medical purposes, such as simulation of possible drug regimens for precision medicine, we expect to broaden our user community. We envisage that by providing the MCA functionality, also users new to mathematical modelling will adapt a working scheme where such sensitivity analyses become an integral part of the model development and study. The value of sensitivity analyses is demonstrated by considering how results of such analyses have given rise to new potential targets for drug discovery [35]. We especially anticipate that the capability of `modelbase` to automatically generate label-specific models will prove useful in predicting fluxes and label propagation dynamics through various metabolic networks. In emerging fields such as computational oncology, such models will be useful to, e.g., predict the appearance of labels in cancer cells. If you have any questions regarding `modelbase`, you are very welcome to ask them. It is our mission to enable reproducible science and to help putting the theory into action.

## Availability and requirements

Project name: `modelbase`

Project home page: https://pypi.org/project/modelbase/

Operating system(s): Platform independent

Programming language: Python

Other requirements: None

Licence: GNU General Public License (GPL), version 3

Any restrictions to use by non-academics: None

## Author’s contributions

MvA developed, tested and maintained the `modelbase` software. OE has conceptualised `modelbase`. AM consulted implemented changes and provided all biomedical examples of `modelbase` implementation. All authors wrote the manuscript.

## Acknowledgements

This work was funded by the Deutsche Forschungsgemeinschaft (DFG, German Research Foundation) under Germany’s Excellence Strategy – EXC-2048/1 – project ID 390686111 (OE, AM), and Deutsche Forschungsgemeinschaft Research Grant MA 8103/1-1 (AM). We would like to thank all members of the Institute of Quantitative and Theoretical Biology for their continuous involvement in the software testing and expansion of the repository of novel and re-implemented computational models.

## Additional Files

### Additional file 1 — Toy model of upper path of glycolysis

Jupyter notebook to reproduce the upper path of glycolysis as introduced in [21].

### Additional file 2 — Labels propagation

Jupyter notebook with an example of a linear non-reversible pathway of 5 randomly sized metabolites and label propagation experiments, as proposed in the paper by Sokol and Portais [23].

### Additional file 3 — Model of the dynamics of HIV infection of CD4^+^T cells

Jupyter notebook with the `modelbase` implementation of the model considering three populations of T cells and free virus, proposed by Perelson, Kirschner and de Boer [29].

### Additional file 4 — Minimal PK-PD model of tumour growth after anticancer therapy

Jupyter notebook with the `modelbase`implementation of the minimal pharmacokinetic-pharmacodynamic (PK-PD)model linking that linking the dosing regimen of an anticancer agent to the tumour growth, proposed by Simeoni and colleagues [34].

### Additional file 5 — Classic epidemic SIR model

Jupyter notebook with the `modelbase` implementation of the classic epidemic Susceptible-Infected-Recovered (SIR) model parametrised as the autonomous model used to simulate periodicity of chicken pox outbreak in Hida, Japan [39].

### Additional file 6 — Official documentations including API changes summary

The official documentation is hosted on ReadTheDocs.

## References

1. Kitano, H.: Computational systems biology. Nature 420(6912), 206–210 (2002). doi:10.1038/nature01254. Number: 6912 Publisher: Nature Publishing Group. Accessed 2020-05-18

2. Barbolosi, D., Ciccolini, J., Lacarelle, B., Barlési, F., André, N.: Computational oncology — mathematical modelling of drug regimens for precision medicine. Nature Reviews Clinical Oncology 13(4) (2016). doi:10.1038/nrclinonc.2015.204. Accessed 2020-05-18

3. Maier, B.F., Brockmann, D.: Effective containment explains subexponential growth in recent confirmed COVID-19 cases in China. Science 368(6492), 742–746 (2020). doi:10.1126/science.abb4557. Publisher: American Association for the Advancement of Science Section: Research Article. Accessed 2020-05-18

4. Tang, B., Bragazzi, N.L., Li, Q., Tang, S., Xiao, Y., Wu, J.: An updated estimation of the risk of transmission of the novel coronavirus (2019-nCov). Infectious Disease Modelling 5, 248–255 (2020). doi:10.1016/j.idm.2020.02.001. Accessed 2020-05-18

5. Rocklöv, J., Sjödin, H., Wilder-Smith, A.: COVID-19 outbreak on the Diamond Princess cruise ship: estimating the epidemic potential and effectiveness of public health countermeasures. J Travel Med 27(3) (2020). doi:10.1093/jtm/taaa030. Publisher: Oxford Academic. Accessed 2020-05-18

6. Brodland, G.W.: How Computational Models Can Help Unlock Biological Systems. ISSN: 1096-3634 Library Catalog: pubmed.ncbi.nlm.nih.gov Publisher: Semin Cell Dev Biol Volume: 47–48 (2015). doi:10.1016/j.semcdb.2015.07.001. https://pubmed.ncbi.nlm.nih.gov/26165820/ Accessed 2020-05-20

7. Butler, J.A., Cosgrove, J., Alden, K., Timmis, J., Coles, M.C.: Model-driven experimentation: A new approach to understand mechanisms of tertiary lymphoid tissue formation, function, and therapeutic resolution. Frontiers in Immunology 8(APR) (2017). doi:10.3389/fimmu.2016.00658

8. Rios-Estepa, R., Lange, I., Lee, J.M., Lange, B.M.: Mathematical Modeling-Guided Evaluation of Biochemical, Developmental, Environmental, and Genotypic Determinants of Essential Oil Composition and Yield in Peppermint Leaves. Plant Physiology 152(April), 2105–2119 (2010). doi:10.1104/pp.109.152256

9. Acs, S., Ostlaender, N., Listorti, G., Hradec, J., Hardy, M., Smits, P., Hordijk, L.: Modelling for EU Policy support: Impact Assessments. Technical report, Publications Office of the European Union, Luxembourg (2019). doi:10.2760/748720 (on-line).

10. Hucka, M., Finney, A., Sauro, H.M., Bolouri, H., Doyle, J.C., Kitano, H., Arkin, A.P., Bornstein, B.J., Bray, D., Cornish-Bowden, A., Cuellar, A.A., Dronov, S., Gilles, E.D., Ginkel, M., Gor, V., Goryanin, I.I., Hedley, W.J., Hodgman, T.C., Hofmeyr, J.-H., Hunter, P.J., Juty, N.S., Kasberger, J.L., Kremling, A., Kummer, U., Le Novère, N., Loew, L.M., Lucio, D., Mendes, P., Minch, E., Mjolsness, E.D., Nakayama, Y., Nelson, M.R., Nielsen, P.F., Sakurada, T., Schaff, J.C., Shapiro, B.E., Shimizu, T.S., Spence, H.D., Stelling, J., Takahashi, K., Tomita, M., Wagner, J., Wang, J.: The systems biology markup language (SBML): a medium for representation and exchange of biochemical network models. Bioinformatics 19(4), 524–531 (2003). doi:10.1093/bioinformatics/btg015. Publisher: Oxford Academic. Accessed 2020-05-20

11. Chelliah, V., Laibe, C., Le Novère, N.: BioModels Database: A Repository of Mathematical Models of Biological Processes. In: Schneider, M.V. (ed.) In Silico Systems Biology. Methods in Molecular Biology, pp. 189–199. Humana Press, Totowa, NJ (2013)

12. Ebenhöh, O., Aalst, M.v., Saadat, N.P., Nies, T., Matuszynśka, A.: Building Mathematical Models of Biological Systems with modelbase. Journal of Open Research Software 6(1), 24 (2018). doi:10.5334/jors.236. Number: 1 Publisher: Ubiquity Press. Accessed 2020-05-18

13. Poolman, M.G.: ScrumPy : metabolic modelling with Python. IEE Proc.-Syst. Biol. 153(5), 375–378 (2006). doi:10.1049/ip-syb

14. Olivier, B.G., Rohwer, J.M., Hofmeyr, J.H.S.: Modelling cellular systems with PySCeS. Bioinformatics 21(4), 560–561 (2005). doi:10.1093/bioinformatics/bti046

15. Schölzel, C., Blesius, V., Ernst, G., Dominik, A.: Required characteristics for modeling languages in systems biology: A software engineering perspective. bioRxiv (April), 2019–1216875260 (2019). doi:10.1101/2019.12.16.875260

16. Andersson, C., Claus, F., Akesson, J.: ScienceDirect Assimulo: A unified framework for ODE solvers. Mathematics and Computers in Simulation 116, 26–43 (2015). doi:10.1016/j.matcom.2015.04.007

17. Hindmarsh, A.C., Brown, P.N., Grant, K.E., Lee, S.L., Serban, R., Shumaker, D.A.N.E., Woodward, C.S.: SUNDIALS : Suite of Nonlinear and Differential / Algebraic Equation Solvers. ACM Transactions on Mathematical Software 31(3), 363–396 (2005)

18. Virtanen, P., Gommers, R., Oliphant, T.E., Haberland, M., Reddy, T., Cournapeau, D., Burovski, E., Peterson, P., Weckesser, W., Bright, J., van der Walt, S.J., Brett, M., Wilson, J., Jarrod Millman, K., Mayorov, N., Nelson, A.R.J., Jones, E., Kern, R., Larson, E., Carey, C., Polat, Ï., Feng, Y., Moore, E.W., Vand erPlas, J., Laxalde, D., Perktold, J., Cimrman, R., Henriksen, I., Quintero, E.A., Harris, C.R., Archibald, A.M., Ribeiro, A.H., Pedregosa, F., van Mulbregt, P., SciPy 1. 0 Contributors: SciPy 1.0: Fundamental Algorithms for Scientific Computing in Python. Nature Methods 17, 261–272 (2020). doi:10.1038/s41592-019-0686-2

19. Kacser, H., Burns, J.A.A.A.: The Control of Flux: 2 1. Symp. Soc. Exp. Biol 27, 65–104 (1973)

20. Heinrich, R., Rapoport, T.A.: A Linear Steady-State Treatment of Enzymatic Chains. General Properties, Control and Effector Strength. European Journal of Biochemistry 42(1), 89–95 (1974). doi:10.1111/j.1432-1033.1974.tb03318.x

21. Klipp, E., Liebermeister, W., Wierling, C., Kowald, A., Lehrach, H., Herwig, R.: Systems Biology: A Textbook. >John Wiley & Sons, ??? (2013)

22. Hunter, J.D.: Matplotlib: A 2d graphics environment. Computing in Science & Engineering 9(3), 90–95 (2007). doi:10.1109/MCSE.2007.55

23. Sokol, S., Portais, J.-C.: Theoretical Basis for Dynamic Label Propagation in Stationary Metabolic Networks under Step and Periodic Inputs. PLoS ONE 10(12), 0144652 (2015). doi:10.1371/journal.pone.0144652. Accessed 2020-05-18

24. Le Novère, N., Finney, A., Hucka, M., Bhalla, U.S., Campagne, F., Collado-Vides, J., Crampin, E.J., Halstead, M., Klipp, E., Mendes, P., Nielsen, P., Sauro, H., Shapiro, B., Snoep, J.L., Spence, H.D., Wanner, B.L.: Minimum information requested in the annotation of biochemical models (MIRIAM). Nature Publishing Group (2005). doi:10.1038/nbt1156

25. Ebenhöh, O., Fucile, G., Finazzi, G., Rochaix, J.-D., Goldschmidt-Clermont, M.: Short-term acclimation of the photosynthetic electron transfer chain to changing light: a mathematical model. Philos. Trans. R. Soc. Lond., B, Biol. Sci. 369(1640), 20130223 (2014). doi:10.1098/rstb.2013.0223

26. Pettersson, G., Ryde-Pettersson, U.: A mathematical model of the Calvin photosynthesis cycle. European Journal of Biochemistry 175(3), 661–672 (1988). doi:10.1111/j.1432-1033.1988.tb14242.x. eprint: https://febs.onlinelibrary.wiley.com/doi/pdf/10.1111/j.1432-1033.1988.tb14242.x. Accessed 2020-05-18

27. McIntyre, L.M., Thorburn, D.R., Bubb, W.A., Kuchel, P.W.: Comparison of computer simulations of the F-type and L-type non-oxidative hexose monophos-phate shunts with 31P-NMR experimental data from human erythrocytes. Eur J Biochem 180(2), 399–420 (1989)

28. Berthon, H.A., Bubb, W.A., Kuchel, P.W.: 13C n.m.r. isotopomer and computer-simulation studies of the non-oxidative pentose phosphate pathway of human erythrocytes. Biochem J 296, 79–387 (1993). doi:10.1042/bj2960379

29. Perelson, A.S., Kirschner, D.E., Boer, R.D.: Dynamics of hiv infection of cd4+ t cells. Mathematical Biosciences 114(1), 81–125 (1993). doi:10.1016/0025-5564(93)90043-A

30. Derendorf, H., Meibohm, B.: Modeling of Pharmacokinetic/Pharmacodynamic (PK/PD) Relationships: Concepts and Perspectives. Pharmaceutical Research 16(2), 176–185 (1999). doi:10.1023/A:1011907920641

31. Lloyd, K.C.K., Khanna, C., Hendricks, W., Trent, J., Kotlikoff, M.: Precision medicine: an opportunity for a paradigm shift in veterinary medicine HHS Public Access. J Am Vet Med Assoc 248(1), 45–48 (2016). doi:10.2460/javma.248.1.45

32. Polasek, T.M., Shakib, S., Rostami-Hodjegan, A.: Precision dosing in clinical medicine: present and future. Taylor and Francis Ltd (2018). doi:10.1080/17512433.2018.1501271

33. Koziol, J.A., Falls, T.J., Schnitzer, J.E.: Different ODE models of tumor growth can deliver similar results. BMC Cancer 20(1), 226 (2020). doi:10.1186/s12885-020-6703-0. Accessed 2020-05-14

34. Simeoni, M., Magni, P., Cammia, C., De Nicolao, G., Croci, V., Pesenti, E., Germani, M., Poggesi, I., Rocchetti, M.: Predictive Pharmacokinetic-Pharmacodynamic Modeling of Tumor Growth Kinetics in Xenograft Models after Administration of Anticancer Agents. Cancer Res 64(3), 1094–1101 (2004). doi:10.1158/0008-5472.CAN-03-2524. Accessed 2020-05-14

35. Cascante, M., Boros, L.G., Comin-Anduix, B., de Atauri, P., Centelles, J.J., W-N Lee, P.: Metabolic control analysis in drug discovery and disease. Nature Biotechnology 20, 243–249 (2002). doi:10.1038/nbt0302-243

36. Hethcote, H.W.: The Mathematics of Infectious Diseases. SIAM Rev. 42(4), 599–653 (2000). doi:10.1137/S0036144500371907

37. Kermack, W.O., McKendrick, A.G.: A contribution to the mathematical theory of epidemics. Proceedings of the Royal Society of London. Series A, Containing Papers of a Mathematical and Physical Character 115(772), 700–721 (1927). doi:10.1098/rspa.1927.0118

38. Wang, J., Liu, S., Zheng, B., Takeuchi, Y.: Qualitative and bifurcation analysis using an SIR model with a saturated treatment function. Mathematical and Computer Modelling 55(3-4), 710–722 (2012). doi:10.1016/j.mcm.2011.08.045

39. Greer, M., Saha, R., Gogliettino, A., Yu, C., Zollo-Venecek, K.: Emergence of oscillations in a simple epidemic model with demographic data. Royal Society Open Science 7(1), 191187 (2020). doi:10.1098/rsos.191187

